# Genetic loci of the *R. anatipestifer* serotype discovered by Pan-GWAS and its application for the development of a multiplex PCR serotyping method

**DOI:** 10.1101/2021.07.26.453894

**Authors:** Zhishuang Yang, Xueqin Yang, Mingshu Wang, Renyong Jia, Shun Chen, Mafeng Liu, Xinxin Zhao, Qiao Yang, Ying Wu, Shaqiu Zhang, Juan Huang, Xumin Ou, Sai Mao, Qun Gao, Di Sun, Bin Tian, Dekang Zhu, Anchun Cheng

## Abstract

The disease caused by *Riemerella anatipestifer* (*R. anatipestifer*) causes large economic losses to the global duck industry every year. Serotype-related genomic variation (such as in O-antigen and capsular polysaccharide gene clusters) has been widely used for the serotyping in many gram-negative bacteria. To date, there have been few studies focused on genetic basis of serotypes in *R. anatipestifer*. Here, we used pan-genome-wide association studies (Pan-GWAS) to identify the serotype-specific genetic loci of 38 *R. anatipestifers* strain. Analyses of the loci of 11 serotypes showed that the loci could be well mapped with the serotypes of the corresponding strains. We constructed the knockout strain for the *wzy* gene at the locus, and the results showed that the mutant lost the agglutination characteristics to positive antisera. Based on the of Pan-GWAS results, we developed a multiple PCR method to identify serotypes 1, 2, and 11 of *R. anatipestifer*. Our study provides a precedent for systematically analysing the genetic basis of the *R anatipestifer* serotypes and establishing a complete serotyping system in the future.

**Highlights:**

1. *R. anatipestifer* serotype-specific locus was identified by Pan-GWAS for the first time.
2. Molecular serotyping multiplex PCR was developed based on O-antigen biosynthesis gene clusters

## Introduction

*R. anatipestifer* attacks domestic ducks, geese, and turkeys and causes an acute or chronic septicaemia characterized by fibrinous pericarditis, perihepatitis, airsacculitis, caseous salpingitis, and meningitis(Boulianne et al.). Since 1982, when Bisgaard(Bisgaard, 1982) established the *R. anatipestifer* serotyping scheme (labelled with Arabic numerals), at least 21 serotypes have been reported around the world(Pathanasophon et al., 2002). There is no effective cross-protection among different serotypes(Liao et al., 2015). Unfortunately, no molecular serotyping method for *R. anatipestifer* has been proposed.

In most gram-negative bacteria, the surface O-antigen structures exhibit intraspecies diversity, which is usually associated with serotypes(Bian et al., 2020; Carter, 1955; Kenyon et al., 2017; Liu et al., 2014; Townsend et al., 2001). The diversity of O-antigens is attributed to genetic variation in O-antigen gene clusters (O-AGCs), which provides a target for molecular serotyping(Fang et al., 2016; Franklin et al., 2011; Liu et al., 2020). Notably, due to the high correlation between the genetic signature of O-AGCs and the serotype phenotype, O-antigen synthesis-related genes (e.g. *wzy*, *wzx* etc.) have been widely used as targets for molecular serotyping of many gram-negative bacteria(Fang et al., 2016; Iguchi et al., 2020; Townsend et al., 2001; Wang et al., 2017a; Xi et al., 2019; Zeng et al., 2019).

Genome-wide association studies (GWAS) have become a powerful tool in bacteria to uncover the genetic basis of some important phenotypes, such as virulence and antibiotic resistance(Farhat et al., 2019; Ma et al., 2020; Young et al., 2019; Yuan et al., 2019; Zankari et al., 2013). In the current study, we used GWAS to identify the genetic loci associated with serotypes and demonstrated that these loci are located within the same genomic region of *R. anatipestifer*. Furthermore, we analysed the genetic diversity of the genetic locus in *R. anatipestifer* (11 serotypes). Based on the genetic variation, we present a multiplex PCR (mPCR) method for the identification of the major serotypes of *R. anatipestifer*, which provides a potential method for epidemiological surveillance of this pathogen.

## Materials and methods

### Bacterial strain

The *R. anatipestifer* strains and the published genome data employed in this study are listed in Supplementary Table 1.

### Agglutination test using the antisera

The serotypes of *R. anatipestifer* involved in this study were determined by slide agglutination according to Brogden et al.(Bisgaard, 1982). Standard typing antisera were purchased from RIPAC-LABOR GmbH (Potsdam, Germany). The *R. anatipestifer* strains were grown on tryptic soy agar (TSA), enriched with 5% sheep blood, at 37°C for 24 h under microaerophilic conditions.

### Genome wide association study of serotypes

To explore the association between *R. anatipestifer* serotypes and genetic characteristics, a pan-genome-wide association study (Pan-GWAS) was performed. Specifically, the *R. anatipestifer* genome was annotated using Prokka v1.12 (Seemann, 2014), and the pan-genome containing 38 strains of *R. anatipestifer* was reconstructed with Roary (Version 3.12.0, with identity threshold of protein = 90)(Page et al., 2015). Furthermore, Scoary(v1.6.16)(Brynildsrud et al., 2016) was used to perform the Pan-GWAS with the *gene_presence_absence* file generated by Roary (only serotypes containing more than 4 strains were considered). Scoary’s P-value and Q-value (P-adjust, adjust algorithm: Benjamini-Hochberg method) cut-offs were set to < 0.05, the sensitivity cut-off was set to 70% and specificity to 85%. Next, we mapped the genes that were significantly associated with the serotype to the corresponding genome to obtain the distribution characteristics. Contig comparisons were generated with Easyfig (v2.2)(Sullivan et al., 2011).

### Functional speculation of the gene cluster

To explore the function of serotype-related genetic loci, genome-wide biosynthetic gene clusters (BGCs) of *R. anatipestifer* was predicted with antiSMASH (version 4.2.0, parameter setting: --clusterblast --subclusterblast --knownclusterblast --smcogs --inclusive --borderpredict) (Blin et al., 2017). AntiSMASH also searches for the most similar gene clusters against the Minimum Information about a Biosynthetic Gene Cluster (MIBiG) database(Medema et al., 2015).

BGCs analysis was performed again by DeepBGC (Hannigan et al., 2019), which uses deep learning strategies to mine biosynthetic gene clusters in the microbial genome. The results of the above two methods will be considered comprehensive.

### Gene boundary determination of *R. anatipestifer* O-AGCs

Based on the results of biosynthetic gene cluster mining, we further determined the boundaries of the *R. anatipestifer* O-antigen gene cluster.

More specifically, we retrieved 509 known O-antigen gene clusters from the NCBI Nucleotide database (https://www.ncbi.nlm.nih.gov/nuccore), which included *Enterobacter*, *Salmonella*, *Yersinia*, etc. (Supplementary Table 2). We downloaded the protein sequence of these gene clusters, used CD-HIT (version 4.8.1, parameter setting: -c 1 -aS 0.95)(Li and Godzik, 2006) to remove redundancies and constructed the O-antigen synthesis gene database. Tblastn was used to map these proteins to the *R. anatipestifer* genome, and the resulting filtering thresholds were as follows: coverage ≥50% (-qcov_hsp_perc 50), e-value≤ 1e-5 (-evalue 1e-5). Subsequently, the densely mapped regions in the genome are considered as candidates for the O-antigen gene cluster. Finally, combined with the prediction results of BGCs, the boundary of the O-antigen gene cluster was determined by manual inspection.

### Annotation of the O-AGCs

Protein-encoding genes were predicted using Prokka v1.12 (Seemann, 2014) with default parameters. To assign functions to the predicted genes, the Conserved Domains Database (CDD)(Marchler-Bauer et al., 2014) was used to search for conserved domains with an E-value threshold of 0.01. Meanwhile characteristic gene annotation of genes was performed using Blastp (v2.6+) against Non-Redundant (NR, https://ftp.ncbi.nlm.nih.gov/blast/db/FASTA/nr.gz). The E-value and query coverage were set at 1e-5 and 50% respectively.

To identify the O-antigen translocase (Wzx) and O-antigen polymerase (Wzy), TMHMM2.0(Krogh et al., 2001) was used to predict the transmembrane regions of proteins.

### Inter- and intra-serotypes comparison of LPS GCs

*wzx*, *wzy* and O-antigen gene cluster nucleotide sequence alignment was performed using MAFFT (Katoh et al., 2002) in automatic mode, and then Mega 7(Kumar et al., 2016) with default parameters and 1000 bootstrap replicates were used to reconstruct the NJ (neighbor joining)(Saitou and Nei, 1987) phylogenetic tree.

Blast (v2.6+) and Easyfig (v2.2)(Sullivan et al., 2011) were used for inter- and intra-serotype O-antigen gene cluster comparisons. DNAMAN (version 9, Lynnon Corp., Quebec, Canada) was used to calculate the percentage homology of protein and DNA sequences.

### Conservation analysis of genetic locus in *Flavobacteriaceae*

To show the conservation of our gene cluster in *Flavobacteriaceae*, the multi-gene search method was implemented. Specifically, Multigeneblast(Medema et al., 2013) was used to find homologues of *R. anatipestifer* O-AGCs from the representative genomes of all species of *Flavobacteriaceae* species. In addition, we used Easyfig (v2.2) to analyse the conservation of the best homologues at the corresponding genus level.

### Development of a multiplex serotyping PCR

Based on the sequence variation in the serotype-specific genes of the O-antigen gene cluster, we designed a primer set with Primer-blast (https://www.ncbi.nlm.nih.gov/tools/primer-blast/index.cgi) and MFEprimer3(Qu and Zhang, 2015), that contains 4 primer pairs to specifically detect each of the 3 *R. anatipestifer* serotypes (Table 2). Three primer pairs Primer_1, Primer_2, and Primer_11 were designed to detect serotypes 1, 2, and 11 respectively. The Primer_RA primer pair serves as an internal control, it can detect all *R. anatipestifer*. Each reaction mixture (25 μl) contained 1 μl template DNA, 12.5 μl Premix Taq (TaKaRa Taq^TM^ Version 2.0 plus dye), Primer_1 (2×0.1 μl), Primer_2 (2×0.2 μl), and Primer_RA (2×0.5 μl).

PCR was conducted with initial denaturation for 5 min at 95°C, followed by 35 cycles of 30 s at 95°C, 30 s at 57.9°C and 1 min at 72°C. PCR products were analysed by agarose gel electrophoresis using 1.5% agarose To evaluate the performance of mPCR, we tested 181 serotype known isolates (n=45, serotype 1; n=79, serotype 2; n=49, serotype 11; n=8, other serotypes) by single-blind method. Cohen’s kappa statistics were performed by R software (version 4.0.3, https://www.r-project.org/) with the package fmsb (version 0.7.0, http://minato.sip21c.org/msb/).

### Construction of *R. anatipestifer wzy* mutant strain CH-2Δ*wzy*

The *wzy* gene (G148_RS04365) was deleted by allelic exchange using the recombinant suicide vector pYA4278 (Kong et al.(Leclercq and Courvalin, 1991); donated by Professor Kong). Briefly, upstream (L) and downstream (R) fragments of the *R. anatipestifer* CH-2 *wzy* gene were amplified by PCR from the genome using *wzy*-Left F and *wzy*-Left R, and *wzy*-Right F and *wzy*-Right R primers, respectively. A 1145-bp Spec^R^ cassette was PCR-amplified from the pYES1 new plasmid using the Spc F and Spc R primers. The three fragments were then spliced together in vitro by overlap extension using the *wzy*-Left F and *wzy*-Right R primers, producing the LSR fragment. Adenosine nucleotides were added to both ends of the PCR product, which was then ligated to the AhdI-digested T-cloning suicide vector pYA4278 to generate pYA4278-LSR, which carries a deletion of the entire *wzy* gene. Subsequently, pYA4278-LSR was successively transformed into *E. coli X7232* and *E. coli* X7213λ pir (Donor) and *R. anatipestifer* CH-2 (Recipient) were mixed in a 10-mM MgSO_4_ solution and incubated on TSB agar with diaminopimelic acid at 37°C for 24 h. Spec^R^ transconjugants were further selected in media containing spectinomycin (40 µg/ml). The detailed steps of this study refer to the methods of Luo et al.(Luo et al., 2015). To confirm the *R. anatipestifer* mutant CH-2Δ*wzy*, we performed PCR targeting the transconjugants (see Figure 12 for details). The primers are listed in Table 2.

## Results

### The serotypes of *R. anatipestifer*

In this study, *R. anatipestifer* involved a total of 11 serotypes, including Serotype 1 (n=7), Serotype 2 (n=12), Serotype 3 (n=1), Serotype 4 (n=1), Serotype 5 (n=1), Serotype 6 (n=3), Serotype 7 (n=3), Serotype 8 (n=1), Serotype 10 (n=5), Serotype 11 (n=4), and Serotype 12 (n=1), which were determined by slide agglutination or from references. All strains and their serotypes are shown in Supplementary Table 1.

### Serotype phenotype of *R. anatipestifer* associated with the gene cluster

To screen for loci associated with serotypes, a GWAS was performed with Scoary on the serotypes containing more than 4 strains (serotypes 1, 2, 10, 11). Under the filtering conditions mentioned in the methods, we obtained a total of 31 target genes. The numbers of genes associated with serotype 1, serotype 2, serotype 10 and serotype 11 were 8, 9, 5 and 8, respectively. The minimum specificity and minimum sensitivity of genes significantly related to the serotype were 85.19% (Serotype 2) and 71.43% (Serotype 1), respectively (Supplementary Table 3). Next, we mapped these genes to the corresponding genome and found that these genes were close to each other and formed a gene cluster. Interestingly, according to the BGCs results predicted by antiSMASH, the gene clusters mentioned above were labelled as lipopolysaccharide biosynthetic gene clusters (Table 1). Thirteen percent of the genes in the gene clusters show similarity with the *Legionella pneumophila* serogroup 1 lipopolysaccharide biosynthesis gene cluster (MIBiG accession:

**Table 1.** Information of lipopolysaccharide biosynthetic gene cluster of *R. anatipestifer* predicted by antiSMASH.

| Serotype | Representative strain | Accession No. of Sequence | Start of Cluster | End of Cluster | Most similar known cluster | MIBiG BGC ID |
| --- | --- | --- | --- | --- | --- | --- |
| 1 | ATCC 11845 | CP003388 | 949593 | 981535 | Lipopolysaccharide biosynthetic gene cluster<br>(13% of genes show similarity);<br>O-antigen biosynthesis gene cluster<br>(10% of genes show similarity) | BGC0000775_c1;<br>BGC0000782_c1 |
| 2 | RA-CH-2 | CP004020 | 911907 | 941443 | Lipopolysaccharide biosynthetic gene cluster<br>(13% of genes show similarity);<br>O-antigen biosynthesis gene cluster<br>(10% of genes show similarity) | BGC0000775_c1;<br>BGC0000782_c1 |
| 10 | RCAD0146 | VLIM00000004 | 135675 | 159655 | Lipopolysaccharide biosynthetic gene cluster<br>(13% of genes show similarity);<br>O-antigen biosynthesis gene cluster<br>(10% of genes show similarity) | BGC0000775_c1;<br>BGC0000782_c1 |
| 11 | RCAD0147 | LUDN01000002 | 137494 | 166627 | Lipopolysaccharide biosynthetic gene cluster<br>(13% of genes show similarity);<br>O-antigen biosynthesis gene cluster<br>(10% of genes show similarity) | BGC0000775_c1;<br>BGC0000782_c1 |

BGC0000775)(Lüneberg et al., 2000; Medema et al., 2015). Ten percent of the genes of the gene clusters show similarity with the *Burkholderia pseudomallei* type II O-antigen biosynthesis gene cluster (MIBiG accession: BGC0000782)(DeShazer et al., 1998). Based on these results, we speculate that the serovar-specific gene cluster was O-antigen biosynthesis gene cluster of *R. anatipestifer*.

We further compared the distribution of the gene cluster between different serotypes, and the results showed that the position of the gene cluster was relatively conserved in the genome of *R. anatipestifer* (Figure 1). In short, the gene region has conserved fragments of 4 and 5 genes (excluding the border) at the beginning and end, respectively (Figure 2).

**Figure 1.**
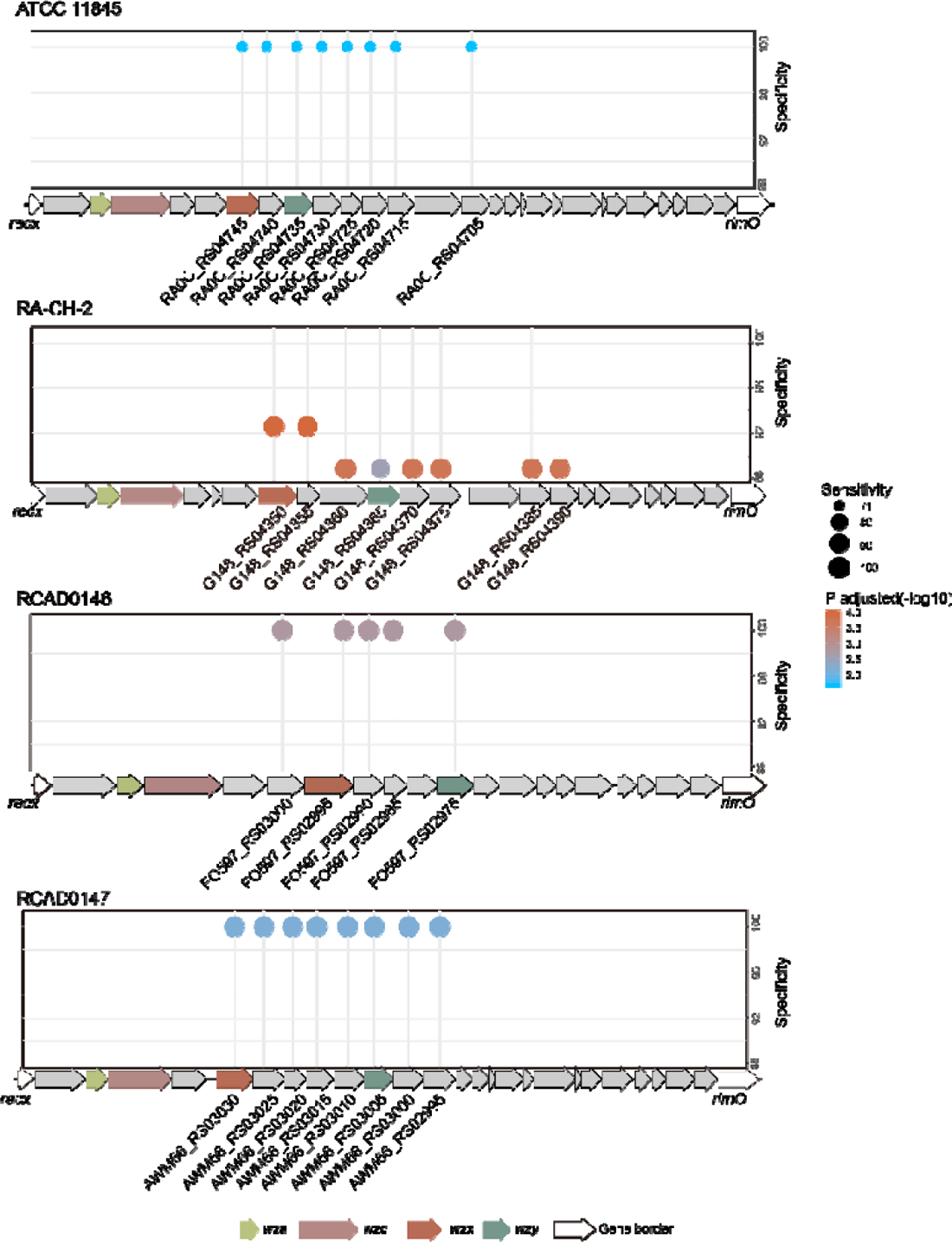
Location of genes significantly associated with serotypes. And the specificity and the sensitivity of genes significantly associated with Serotype 1, 2, 10 and 11. The size of the shape indicates sensitivity; colour indicates negative value of *P*-value (adjusted).

**Figure 2.**
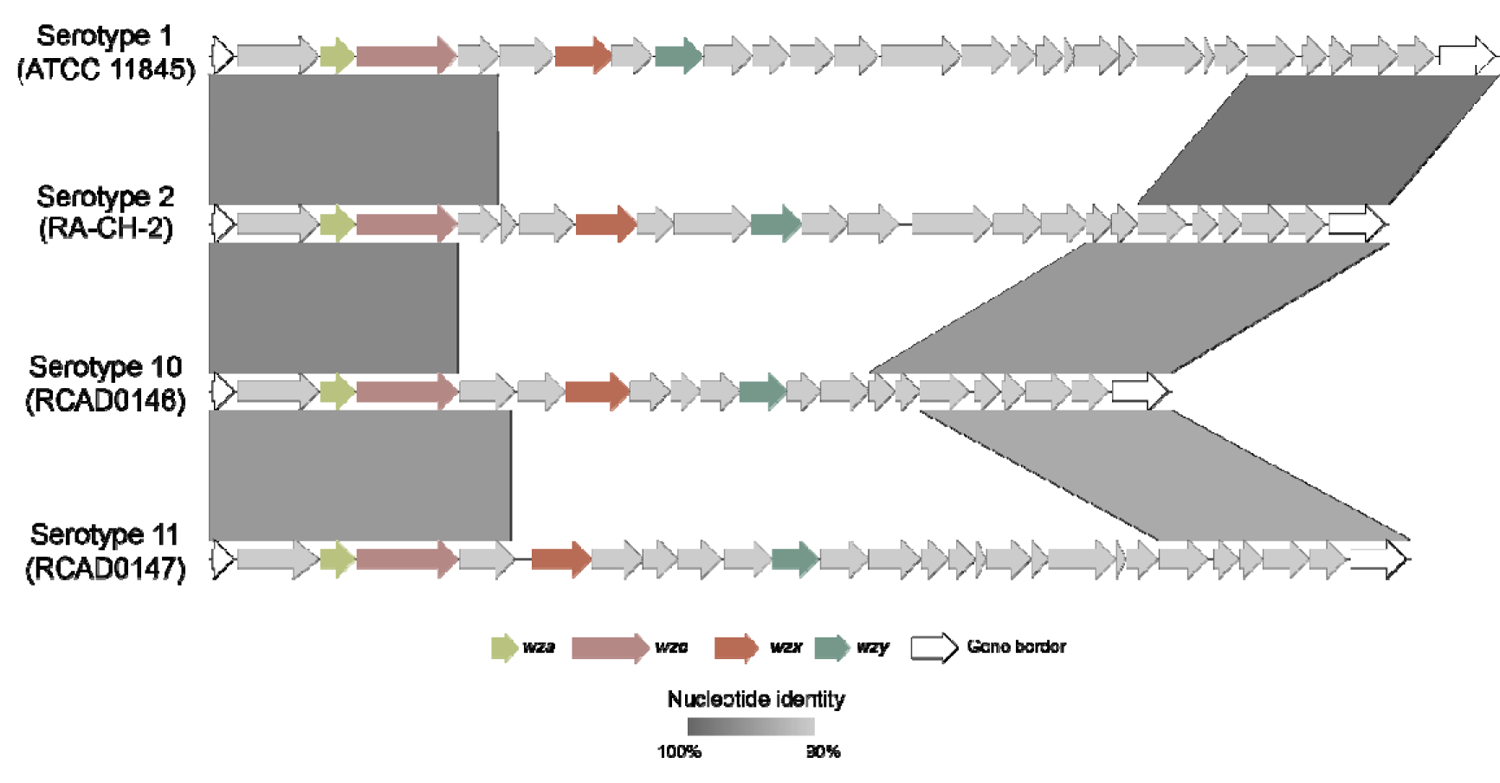
Comparison of serotype-associated regions.

### Analysis of O-antigen gene cluster

To determine the boundaries of the O-antigen gene cluster, we focused on the locations where those O-antigen synthesis genes were densely located. Meanwhile, we reviewed the aforementioned lipopolysaccharide biosynthetic gene clusters, which were conserved in *R. anatipestifer*. Both antiSMASH and DeepBGC characterize a BGC at positions 908750--941443 (RA-CH-2, CP004020.1). According to the antiSMASH results, 10% of the genes in this BGC and the O-antigen gene cluster (GenBank accession: AF064070.1) show similarity. Interestingly, there is a dividing line around the distribution of genes involved in O-antigen synthesis near this area (Figure 3). Several other serotypes had the same situation (Supplementary Figure 1).

**Figure 3.**
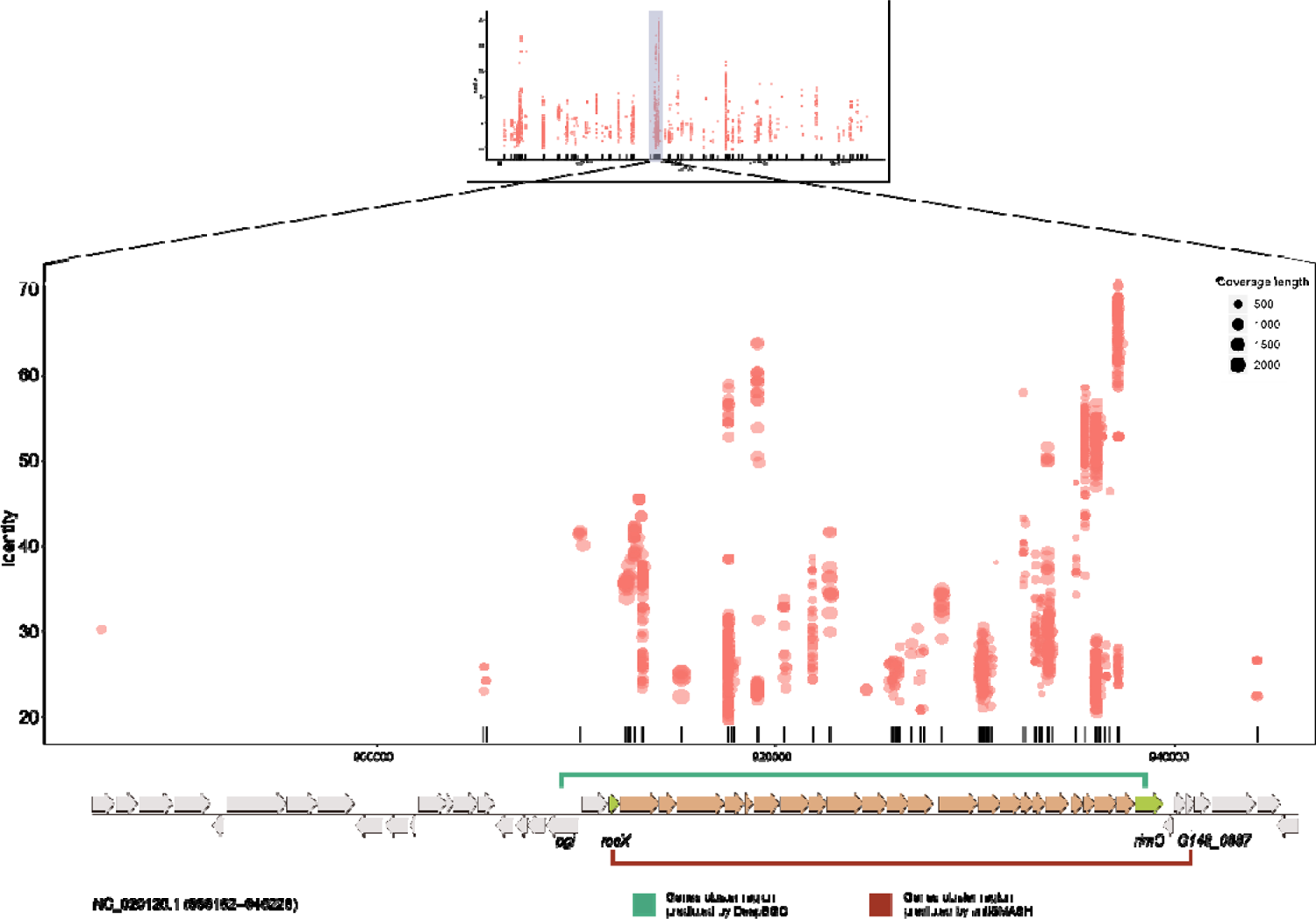
CH-2 O-antigen gene cluster location and boundary determination. The dot plot represents the hits of genes related to O-antigen synthesis on the genome, and the size of the dot indicates the coverage length. Interval markers on gene clusters indicate the BGC regions predicted by DeepBGC and antiSMASH.

Therefore, we speculated that the O-antigen gene cluster of RA-CH-2 was located between *recX* (recombinase, Accession No. G148_RS04315) and *rimO* (ribosomal protein S12 methylthiotransferase, Accession No. G148_RS04430), both of which were highly conserved in *R. anatipestifer*.

The O-antigen gene cluster of RA-CH-2 was 25.6 kb, and the G + C content was 34.00%. It included 22 open reading frames (ORFs) with the same transcriptional direction (Figure 5). To assign annotations the genes, BLAST searches against the NR database and CDD were performed (Table 3).

**Table 2.**
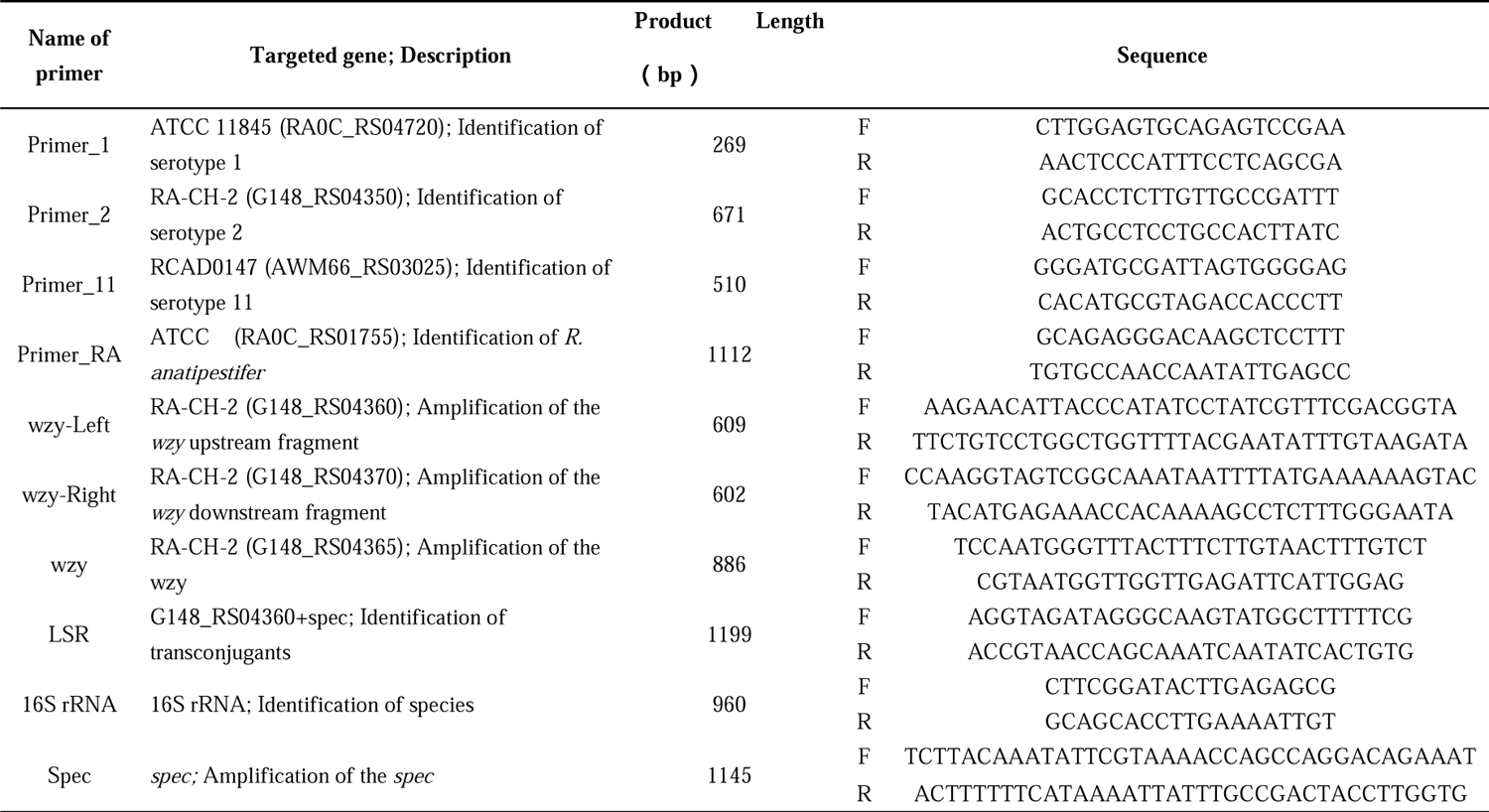
Oligonucleotide primers used in this study

**Table 3.**
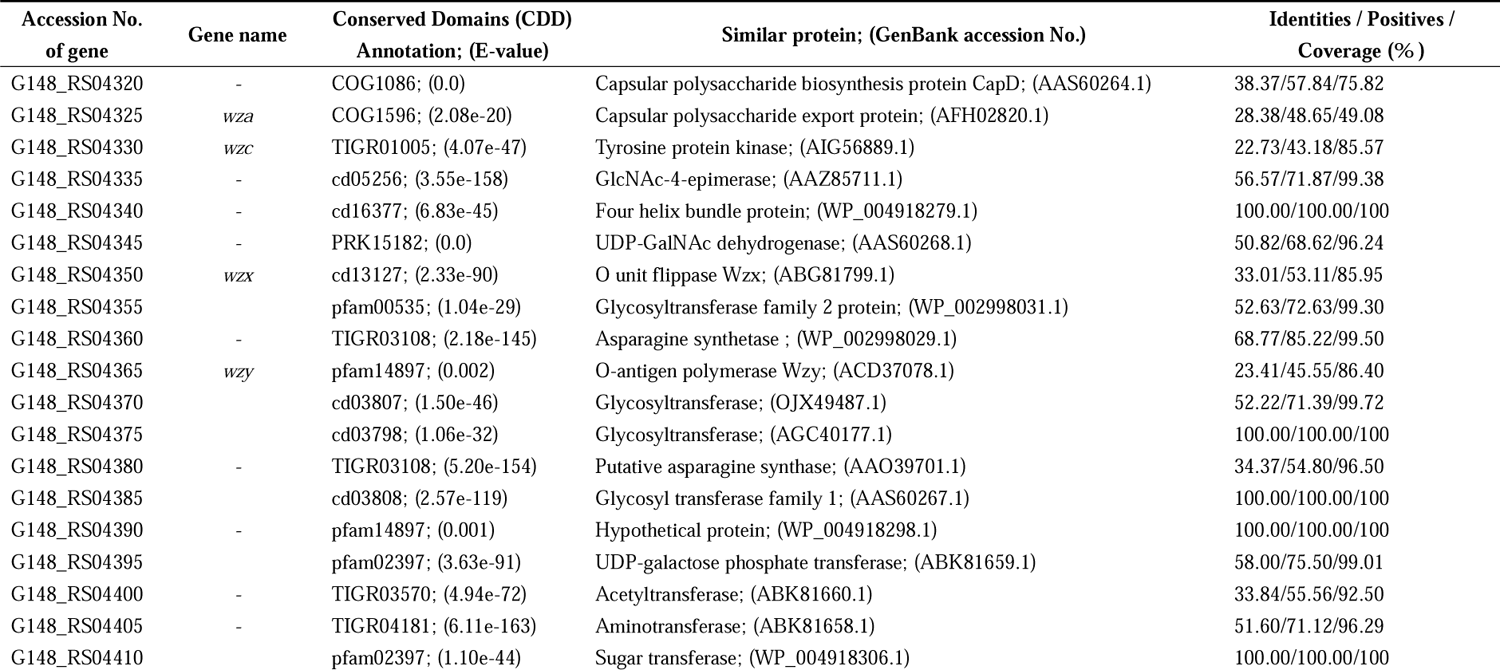
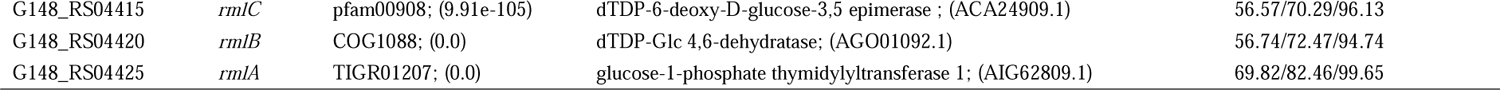
Functional prediction of the gene in the O-antigen gene cluster of *R. anatipestifer* (CH-2, serotype 2)

| Accession No.<br>of gene | Gene name | Conserved Domains (CDD)<br>Annotation; (E-value) | Similar protein; (GenBank accession No.) | Identities / Positives /<br>Coverage (%) |
| --- | --- | --- | --- | --- |
| G148_RS04320 | - | COG1086; (0.0) | Capsular polysaccharide biosynthesis protein CapD; (AAS60264.1) | 38.37/57.84/75.82 |
| G148_RS04325 | wza | COG1596; (2.08e-20) | Capsular polysaccharide export protein; (AFH02820.1) | 28.38/48.65/49.08 |
| G148_RS04330 | wzc | TIGR01005; (4.07e-47) | Tyrosine protein kinase; (AIG56889.1) | 22.73/43.18/85.57 |
| G148_RS04335 | - | cd05256; (3.55e-158) | GlcNAc-4-epimerase; (AAZ85711.1) | 56.57/71.87/99.38 |
| G148_RS04340 | - | cd16377; (6.83e-45) | Four helix bundle protein; (WP_004918279.1) | 100.00/100.00/100 |
| G148_RS04345 | - | PRK15182; (0.0) | UDP-GalNAc dehydrogenase; (AAS60268.1) | 50.82/68.62/96.24 |
| G148_RS04350 | wzx | cd13127; (2.33e-90) | O unit flippase Wzx; (ABG81799.1) | 33.01/53.11/85.95 |
| G148_RS04355 | - | pfam00535; (1.04e-29) | Glycosyltransferase family 2 protein; (WP_002998031.1) | 52.63/72.63/99.30 |
| G148_RS04360 | - | TIGR03108; (2.18e-145) | Asparagine synthetase ; (WP_002998029.1) | 68.77/85.22/99.50 |
| G148_RS04365 | wzy | pfam14897; (0.002) | O-antigen polymerase Wzy; (ACD37078.1) | 23.41/45.55/86.40 |
| G148_RS04370 | - | cd03807; (1.50e-46) | Glycosyltransferase; (OJX49487.1) | 52.22/71.39/99.72 |
| G148_RS04375 | - | cd03798; (1.06e-32) | Glycosyltransferase; (AGC40177.1) | 100.00/100.00/100 |
| G148_RS04380 | - | TIGR03108; (5.20e-154) | Putative asparagine synthase; (AAO39701.1) | 34.37/54.80/96.50 |
| G148_RS04385 | - | cd03808; (2.57e-119) | Glycosyl transferase family 1; (AAS60267.1) | 100.00/100.00/100 |
| G148_RS04390 | - | pfam14897; (0.001) | Hypothetical protein; (WP_004918298.1) | 100.00/100.00/100 |
| G148_RS04395 | - | pfam02397; (3.63e-91) | UDP-galactose phosphate transferase; (ABK81659.1) | 58.00/75.50/99.01 |
| G148_RS04400 | - | TIGR03570; (4.94e-72) | Acetyltransferase; (ABK81660.1) | 33.84/55.56/92.50 |
| G148_RS04405 | - | TIGR04181; (6.11e-163) | Aminotransferase; (ABK81658.1) | 51.60/71.12/96.29 |
| G148_RS04410 | - | pfam02397; (1.10e-44) | Sugar transferase; (WP_004918306.1) | 100.00/100.00/100 |
| G148_RS04415 | *rmlC* | pfam00908; (9.91e-105) | dTDP-6-deoxy-D-glucose-3,5 epimerase ; (ACA24909.1) | 56.57/70.29/96.13 |
| G148_RS04420 | *rmlB* | COG1088; (0.0) | dTDP-Glc 4,6-dehydratase; (AGO01092.1) | 56.74/72.47/94.74 |
| G148_RS04425 | *rmlA* | TIGR01207; (0.0) | glucose-1-phosphate thymidyltransferase 1; (AIG62809.1) | 69.82/82.46/99.65 |

Generally, the coding sequence within O-antigen gene clusters primarily consists of the following three categories: nucleotide sugar biosynthesis, glycosyl transferase, and O-antigen processing(Kalynych et al., 2014). Among gram-negative bacteria, O-antigen processing enzymes include a flippase (Wzx) and O-antigen polymerase (Wzy), which are involved in the transmembrane transport of O-units and the synthesis of O-antigens, respectively(Kalynych et al., 2014). It is worth noting that *wzx* and *wzy* are usually used as serotype molecular detection targets due to their serotype specificity.

To analyse the oligosaccharide unit processing genes, we used TMHMM2.0 to predict the transmembrane domains in the proteins. The results showed that *G148_RS04350* and *G148_RS04365* contain multiple transmembrane regions (Figure 4). A 50 amino acid stem-loop structure is located between the second and third transmembrane regions (Figure 4 b).The large stem-loop structure distributed in the periplasmic region is a typical feature of the O-antigen polymerase (Wzy)(Daniels et al., 1998). Furthermore, the protein of *G148_RS04365* shared 23.41% identity, 45.55% similarity and 86.40% coverage with Wzy (ACD37078.1) from *Shigella boydii* (Table 3).

**Figure 4.**
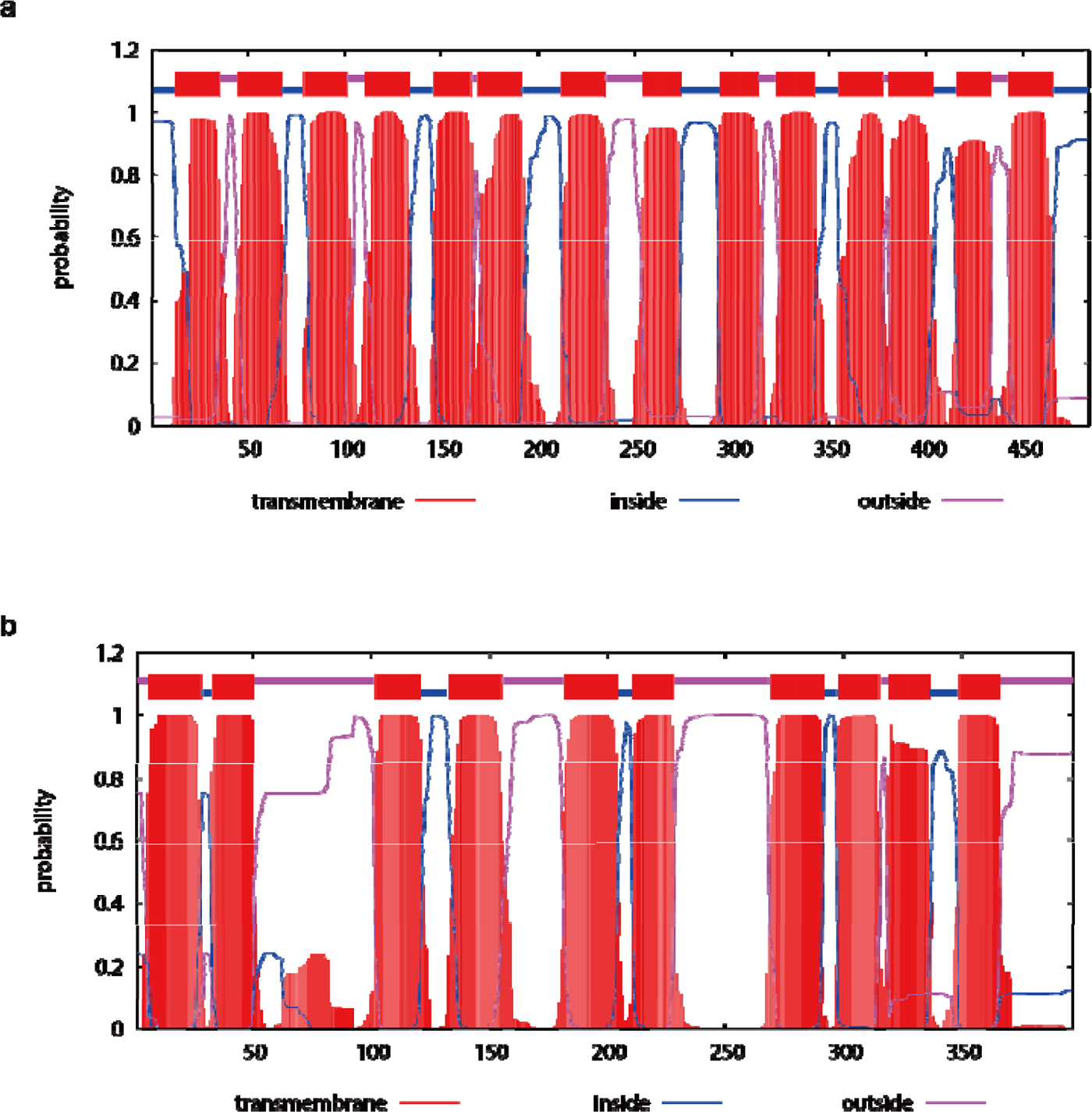
The prediction of transmembrane helices in amino acid sequences encoded by *wzx* and *wzy*. a) The prediction of transmembrane helices in the amino acid sequence encoded by *G148_RS04350*. b) The prediction of transmembrane helices in the amino acid sequence encoded by *G148_RS04365*.

The protein encoded by *G148_RS04350* contains 14 uniformly distributed transmembrane regions (Figure 4 a). *G148_RS04350* shows 33.01% identity, 53.11% similarity and 85.95% coverage to the O-unit flippase (ABG81799.1, AJR19423.1) in *E. coli* and 32.67% identity, 52.32% similarity and 92.35% coverage to the O-unit flippase in *Providencia alcalifaciens*. The set of *wzx* and *wzy* genes suggests the presence of Wzx/Wzy pathway related O-antigen processing.

### Inter- and intra-serotypes comparison of O-AGCs

Based on the positional conservation of the O-antigen gene cluster, we extracted the O-antigen gene cluster sequences from other strains (34 strains Supplementary Table 4). The length of the gene clusters from 20.22 kb (RCAD0135) to 28.38 kb (RCAD0179), GC content between 32.26% (CCUG25011) and 34.00% (RCAD0123), which was significantly lower than the GC content of the genome (upper quartile: 35.05%, lower quartile: 34.97%, mean: 35.00%; Wilcoxon test: p-value = 5.476e-16). These gene clusters contain an average of 25 CDSs (ranging from 20 to 30). We annotated the O-antigen gene clusters of the serotype representative strains marked in Supplementary Table 1, the results are shown in Supplementary Table 5 and Figure 5. It is worth noting that all serotype O-antigen gene clusters contain *wza*, *wzc* and *rmlABC* homologous genes. *rmlABCD* in the serotype 12 gene cluster implies the possible presence of rhamnose in O-units. The set of *wzx* and *wzy* genes indicates the presence of Wzx/Wzy-related O-antigen processing pathways in the corresponding serotypes.

**Figure 5.**
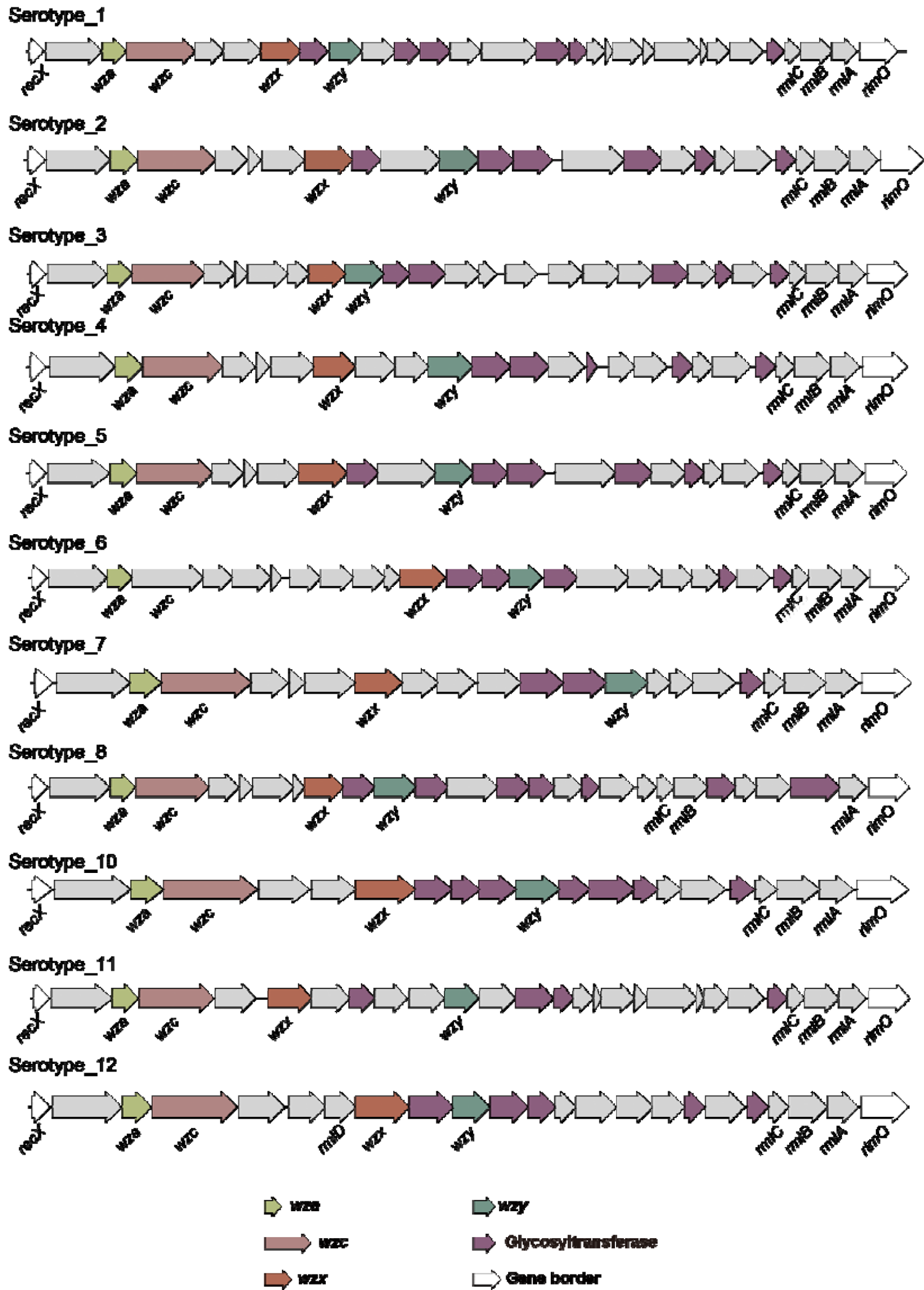
The O-antigen gene cluster in *R. anatipestifer* serotypes 1, 3, 4, 5, 6, 7, 9, 10, 11 and 12

Next, we extracted the *wzx* and *wzy* gene sequences of all the strains and constructed the NJ phylogenetic tree (Figure 6). The homology matrix of the protein or DNA sequences of the *wxy*, *wzx*, and O-antigen gene clusters was calculated, and the results are shown in Figure 7. Overall, *wzx* and *wzy* are serotype-specific, and much greater differences exist among the different serotypes. Interestingly, RA-YM and RA-GD are reported to be serotype 1(Yuan et al., 2011; Zhou et al., 2011), but their *wzx* and *wzy* are highly identical to our reported serotype 2. CCUG25011 is serotype 4, and *wzx* shares more than 97% similarity with serotype 7. Similarly, CCUG25004 is serotype 5, but *wzy* shares high homology with serotype 2(similarity > 99 %).

**Figure 6.**
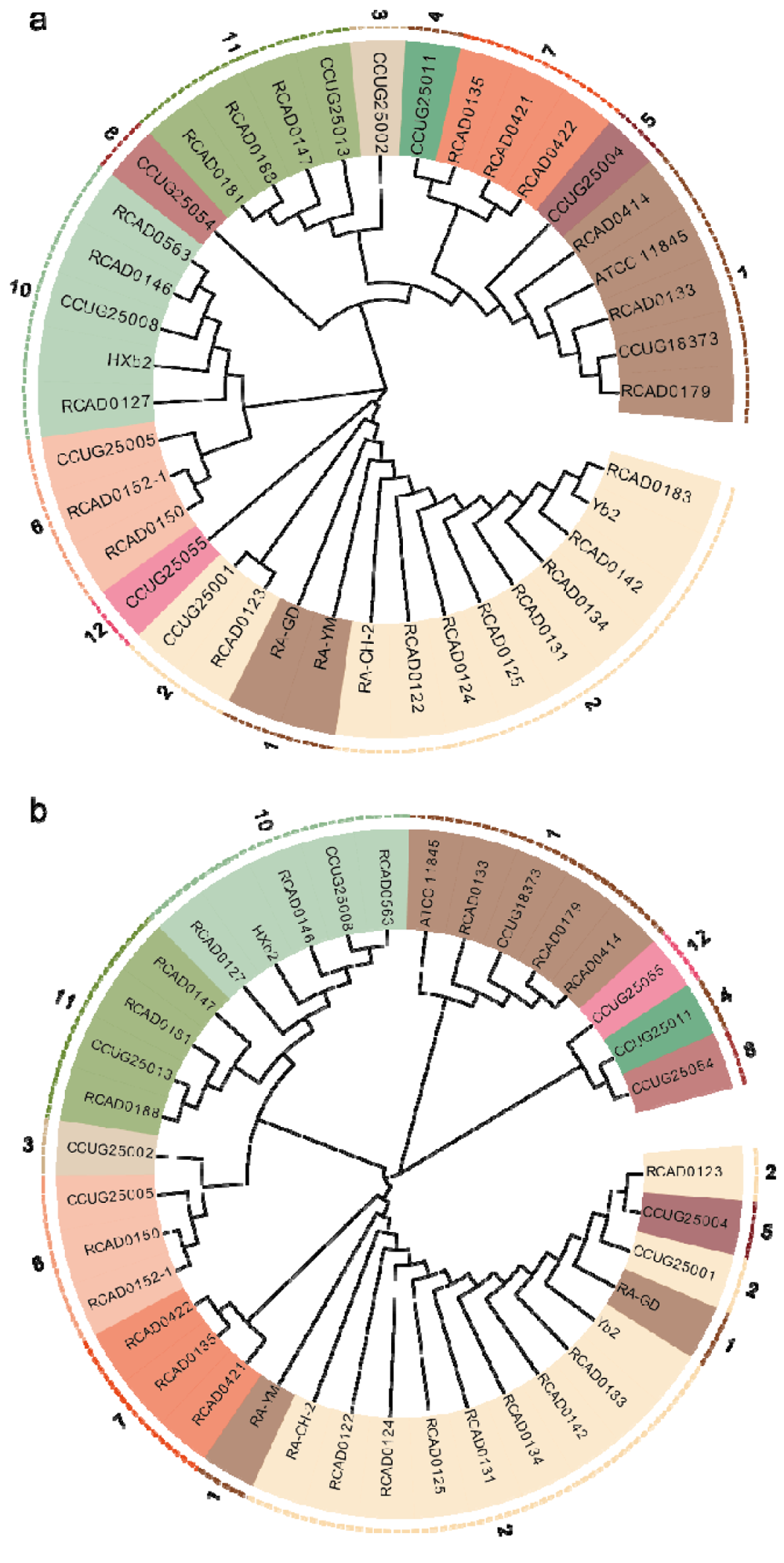
Phylogenetic tree constructed by the neighbor joining method based on the *wzx* (a) and *wzy* (b) genes. Numbers in the outer circle indicate serotypes.

**Figure 7.**
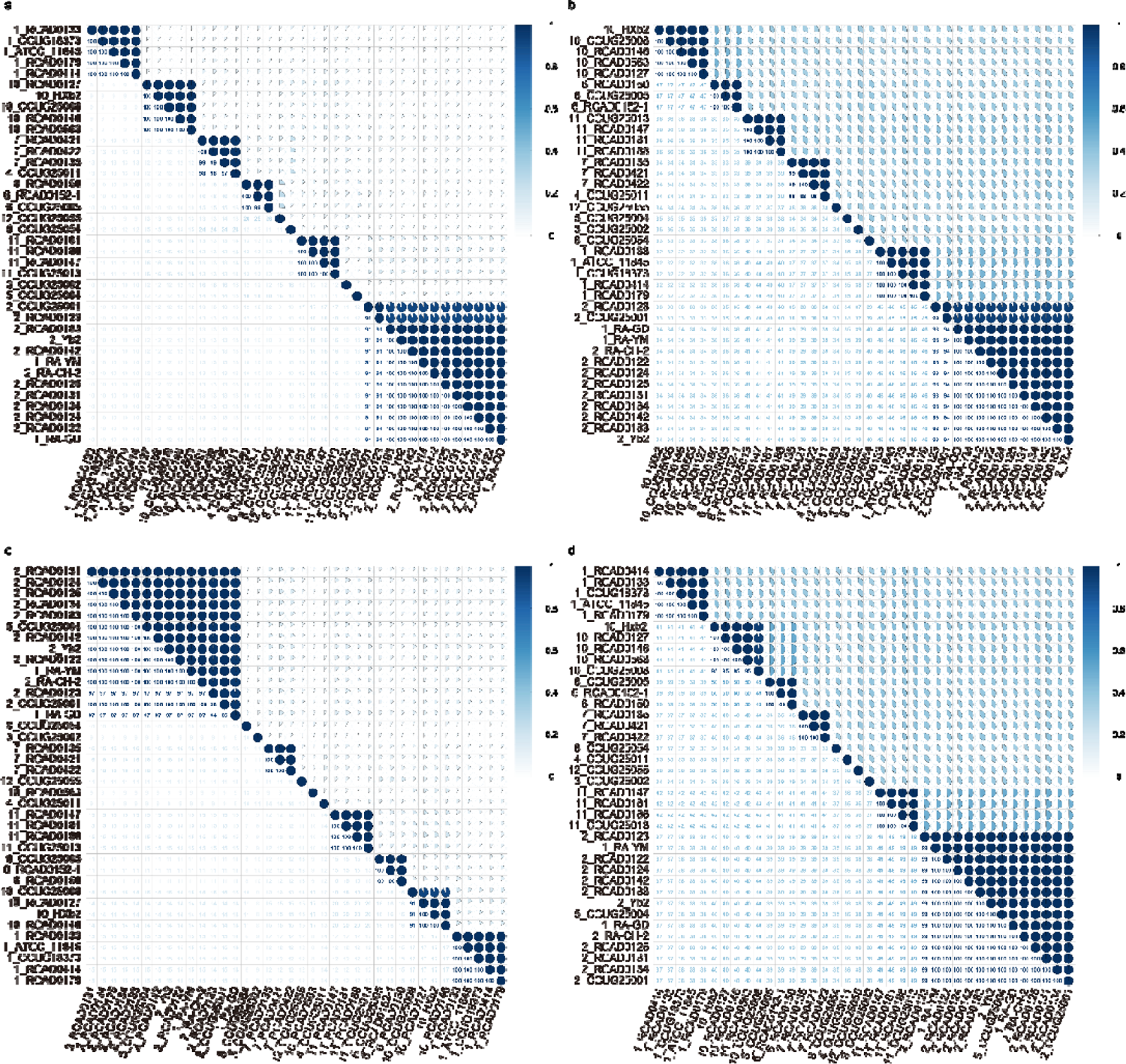
*wzx* and *wzy* homology matrix calculated by DNAMAN. The value in front of the strain ID indicates the serotype. a) Protein homology matrix encoded by the *wzx* gene. b) Homology matrix of the *wzx* gene. c) Protein homology matrix encoded by the *wzy* gene. d) Homology matrix of the *wzy* gene.

In addition, an NJ phylogenetic tree based on the complete sequence of the O-antigen gene cluster and a synteny analysis of the O-antigen gene cluster (DNA sequence identity cut-off: 69%) are shown in Figure 8. As expected, strains of the same serotype clustered into the same clade and had the same gene cluster structure. Consistent with the single gene identification results, the O-antigen gene clusters of RA-YM and RA-GD are highly homologous to the serotype 2 strain. The *wzy* gene of RA-GD is divided into *wzy1* (RIA_1497) and *wzy2* (RIA_1498) according to GenBank. However, the remaining serotype 1 strains were clustered in other clades and had a consistent genetic structure with each other. Except for a gene insertion event in CCUG25001, the genetic structure of the O-antigen gene cluster of all serotype 2 strains was highly similar (Figure 8). The inserted gene predictive function is O-acetylase involved in peptidoglycan or LPS synthesis (Reference: RBP22008.1; Identity/coverage: 40.95%/99%; E-value: 3e-59). Additionally, CCUG25004 (serotype 5) and the strains of serotype 2 differed by only two genes (Wzx and a glycosyltransferase, Figure 8 and Supplementary Table 5). As expected, serotype 1 strains had identical gene clusters. In contrast to serotype 1, the gene clusters among the serotype 10 strains are more diverse, and even so, their *wxz* and *wxy* are uniform. RCAD0127 and HXb2 have almost the same sequence. Gene clusters in serotype 4 and serotype 7 have high identity, *wzx* is a homologue, but *wxy* is specific (Supplementary Figure 2). Overall, serotype 11 shows reasonable sequence homology.

**Figure 8.**
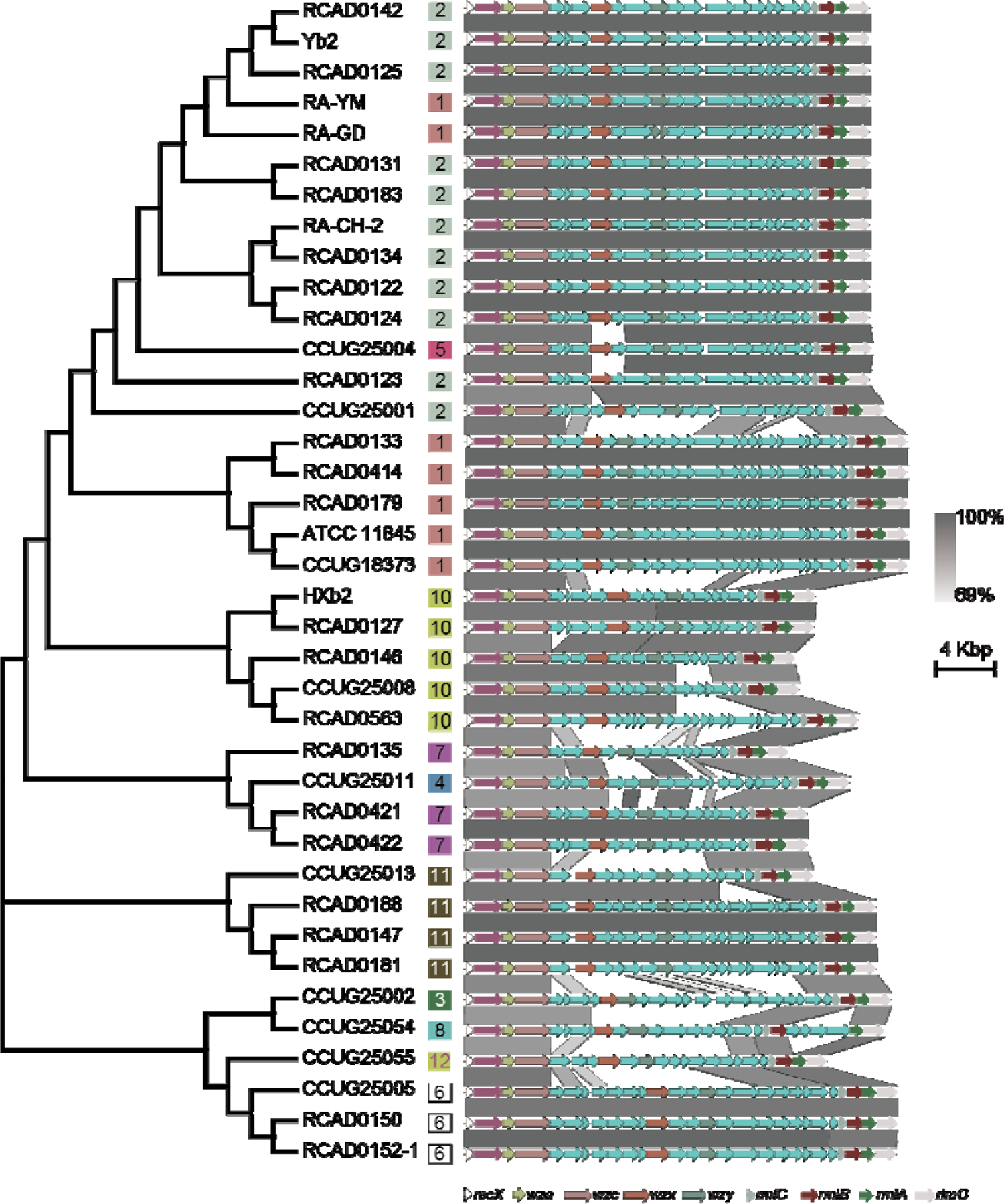
Neighbor-joining phylogenetic tree and structure of the O-antigen gene cluster. Numbers in boxes indicate serotypes

### Conserved loci in other *Flavobacteriaceae* species

The synteny analysis of homologous gene clusters in *Flavobacteriaceae* indicated that the locus of the O-AGC locus was conserved among closely related species (Figure 9a, Supplementary Figure 3). Specifically, the upstream gene arrangement (*recx*-*gdr*-*wza*-*wzc*) of *R. anatipestifer* O-AGCs was highly conserved. *Chryseobacterium* and *R. anatipestifer* were the same (*recx* and *rimO*) at the beginning and end of the region.

**Figure 9.**
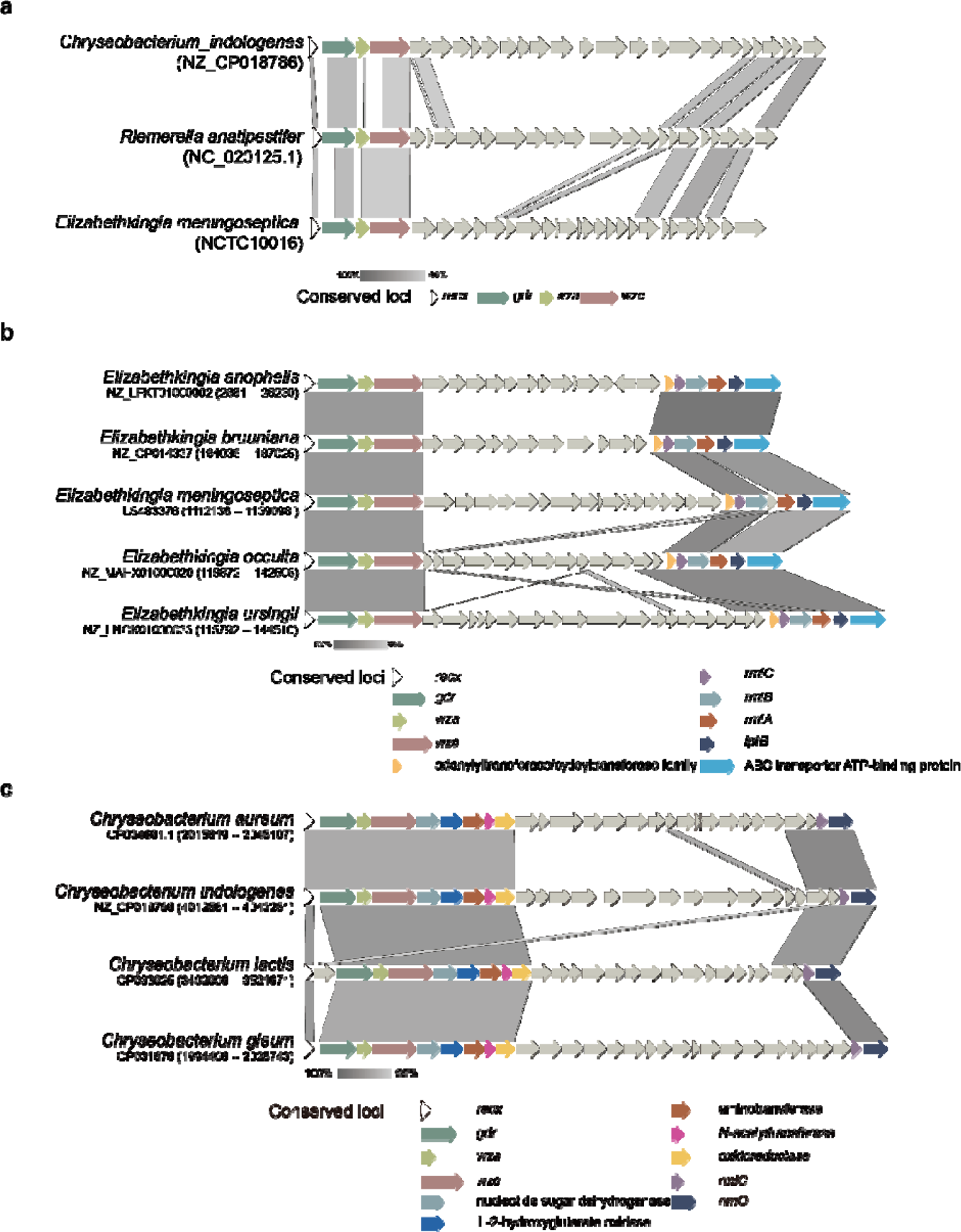
Conserved loci in other *Flavobacteriaceae* species. a) The genetic locus of the O-antigen gene cluster in *R. anatipestifer* is conserved among the closest species. b) Conserved structure in multiple *Elizabethkingia* species. c) Conserved structure in multiple *Chryseobacterium* species.

As expected, this locus is also conserved in *Elizabethkingia sp.* And *Chryseobacterium sp.* (Figure 9b and Figure 9c). Furthermore, many glycosyltransferases related to polysaccharide synthesis are distributed in this region in both genera. It is worth mentioning that *rmlABC* (*Elizabethkingia sp.*), lipopolysaccharide export system ATP-binding protein gene (*lptB, Elizabethkingia sp.*), O-antigen ligase gene (*Chryseobacterium sp.*), and oligosaccharide flippase gene (*Chryseobacterium sp.*) were also present in the conserved region, and they are usually involved in the synthesis of O-antigen or lipopolysaccharide.

Regarding the other two species of *Riemerella*: *Riemerella columbina* and *Riemerella columbipharyngis*, a similar gene cluster was found in *Riemerella columbina* DSM 16469 (Supplementary Figure 4). Furthermore, the genes encoding oligosaccharide repeat unit polymerase (Wzy) and oligosaccharide flippase (Wzx) were annotated in the cluster. However, compared with *Riemerella anatipestifer* RA-CH-2 O-AGCs, the cluster region is significantly rearranged in *Riemerella columbina*. Unfortunately, due to a lack of data, we could not detect similar genetic regions in *Riemerella columbipharyngis*.

### Multiplex serotyping PCR of *R. anatipestifer* serotype 1, 2 and 11

Based on comparative analysis of the inter- and intra-serotypes, we identified specific sites for the three major serotypes. A multiplex PCR method was developed for molecular serotyping (Table 2 and Figure 10). For *R. anatipestifer* serotypes 1,2 and 11, serotyping PCR can produce bands of the correct size for serotyping and species identification (Figure 10a). For the other serotypes (3, 4, 5, 6, 7, 8, 10 and 12), PCR only amplified the species’s identification band (Figure 10c). Moreover, our method has high specificity for common avian pathogens. Even *Riemerella columbina*, belongs to the same genus as *R. anatipestifer* (Figure 10b).

**Figure 10.**
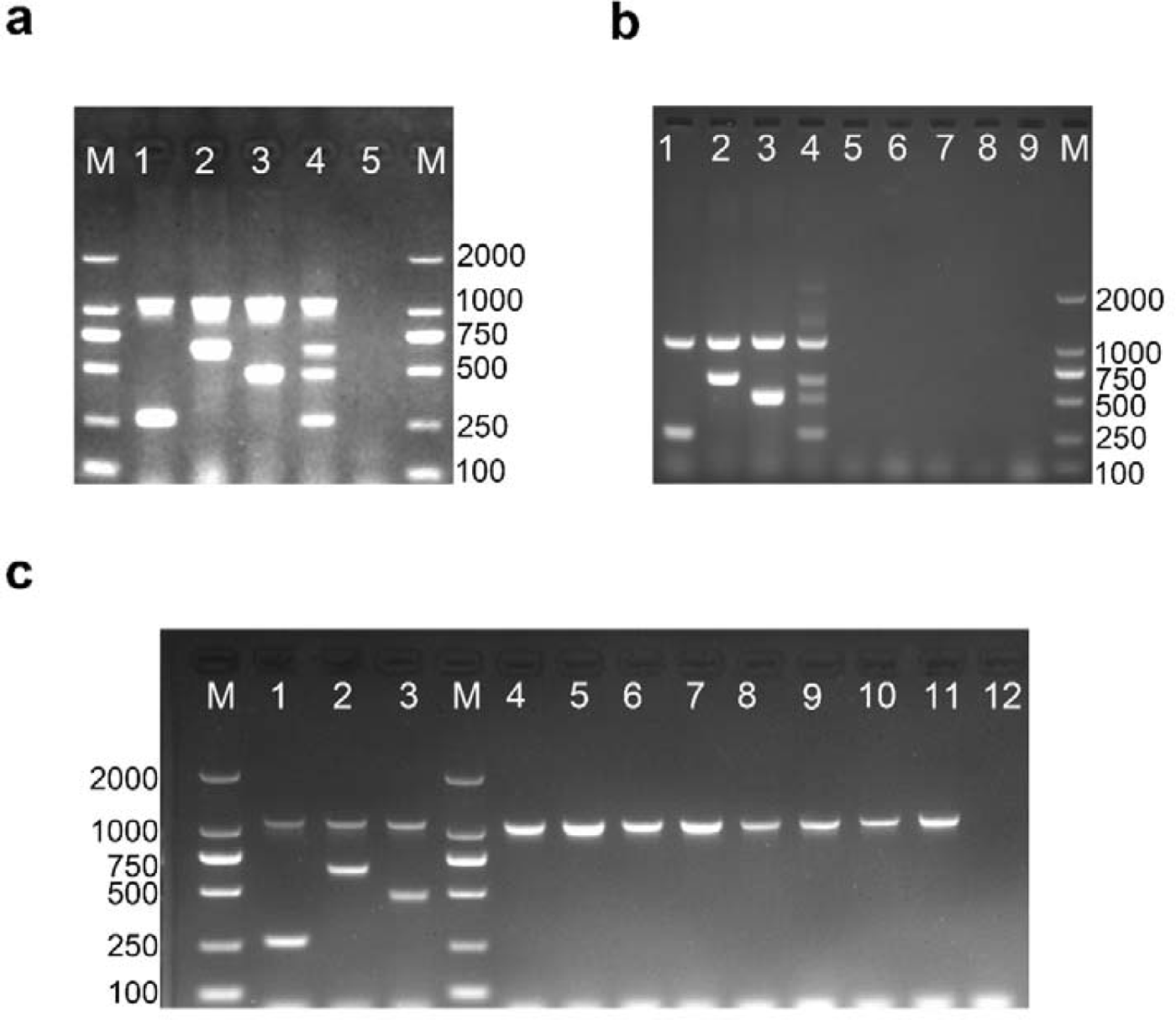
Multiplex PCR method for the identification of *R. anatipestifer* serotypes 1,2 and 11. Lane M, 2000 bp DNA ladder; a) Multiplex PCR method for the identification of *R. anatipestifer* serotypes 1,2 and 11: Lane 1: serotype 1 (ATCC 11845); lane 2: serotype 2 (RA-CH-2); lane 3: serotype 11 (RCAD0147); lane 4: mixed serotypes (1,2,11); and lane 5: Negative control. b) Detection of species specificity for the multiplex PCR method. Lanes 1 to 4: serotype 1 (ATCC 11845), serotype 2 (RA-CH-2), serotype 11 (RCAD0147), and mixed serotypes (1,2,11); lanes 5 to 8: *Pasteurella multocida*, *Salmonella enterica*, *Riemerella columbina*, *Escherichia coli*; and lane 9: negative control. c) Detection of serotype specificity for the multiplex PCR method. Lanes 1 to 3: serotype 1 (ATCC 11845), serotype 2 (RA-CH-2) and serotype 11 (RCAD0147); lanes 4 to 11: serotype 3 (CCUG25002), serotype 4 (CCUG25011), serotype 5 (CCUG25004), serotype 6 (CCUG25005), serotype 7 (CCUG25010), serotype 8 (CCUG25054), serotype 10 (RCAD0146) and serotype 12 (CCUG25055); and lane 12: negative control.

**Figure 11.**
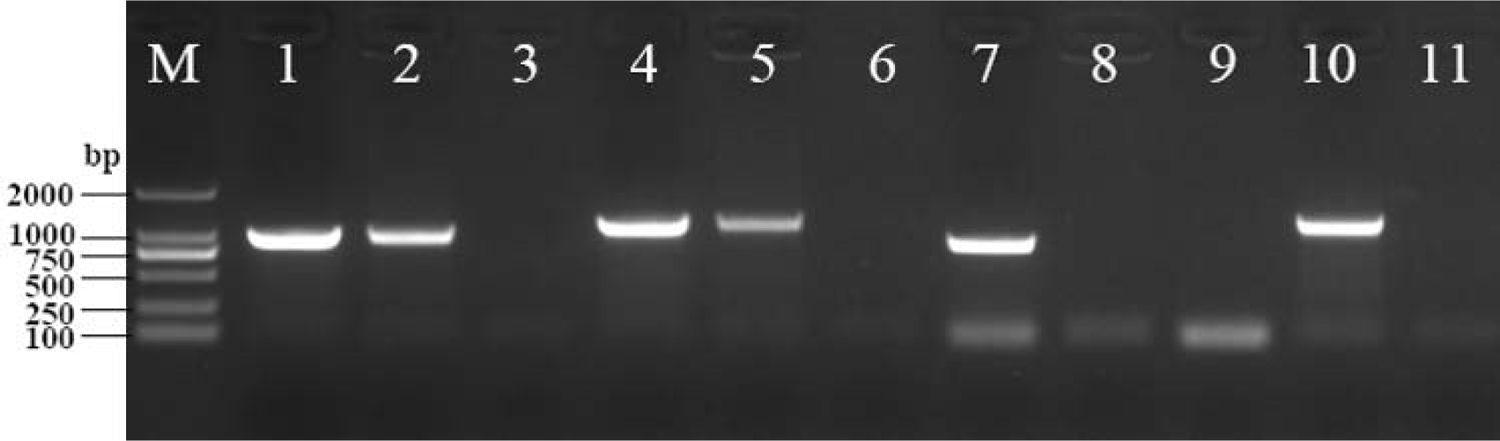
Identification of *R. anatipestifer* CH-2Δ wzy. M: DL2000 DNA Marker; Lanes 1-3: 16S rRNA F and 16S rRNA R, which amplify a 960 bp fragment from *R. anatipestifer* 16S rRNA. Order: Wild-type(CH-2), mutant(CH-2Δ negative control (distilled water); Lanes 4-6: Spec F and Spec R, which amplify a 1180 bp fragment from the SpecR cassette. Order: Positive control (pYES1 new), mutant(CH-2Δ wzy), and negative control (distilled water); Lanes 7-9: wzy F and wzy R, which amplify an 886 bp fragment from the wzy gene. Order: Wild-type(CH-2), mutant(CH-2Δ *wzy*), and negative control (distilled water); Lanes 10-11: LSR F and LSR R, which amplify a 1199 bp fragment from the SpecR cassette, indicating that it was inserted in the correct position in the *R. anatipestifer* CH-2 genome. Order: Mutant(CH-2Δ*wzy*), Negative control (distilled water).

**Figure 12.**
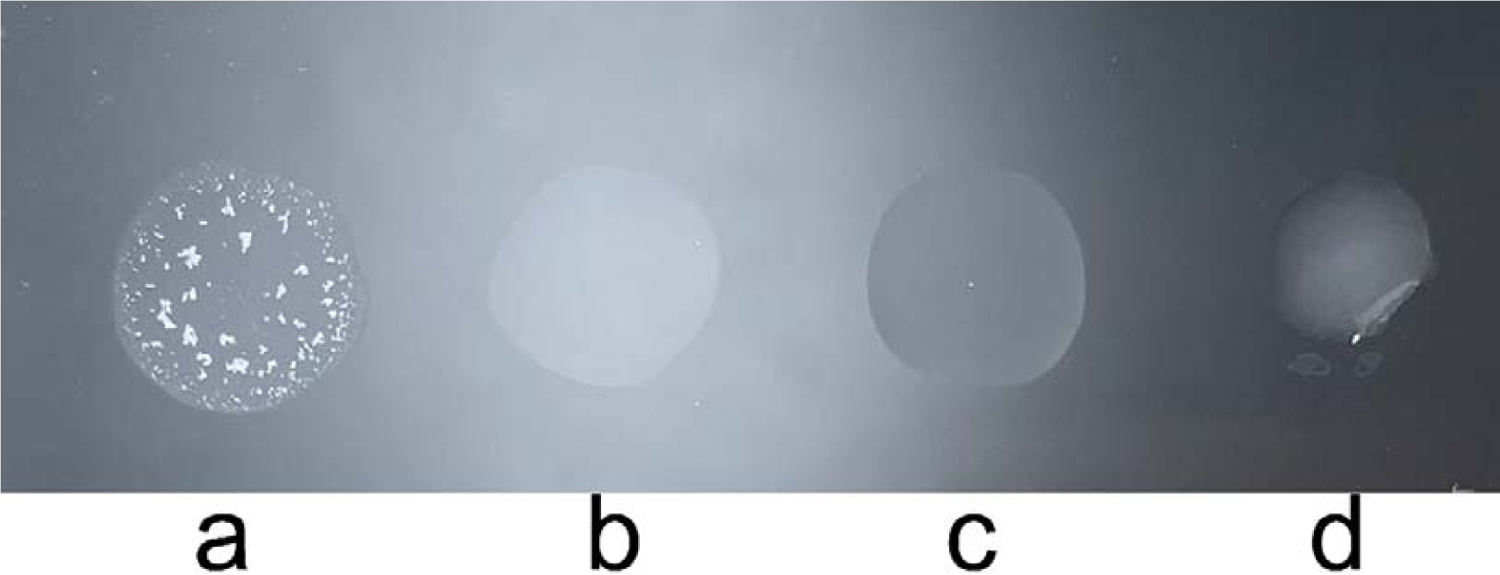
Agglutination test of *R. anatipestifer* CH-2Δ*wzy*. a) Wild-type(CH-2) suspension mixed with antisera of serotype 2; b) mutant(CH-2Δ mixed with antisera of serotype 2; c) distilled water mixed with antisera of serotype 2; and d) mutant(CH-2Δ*wzy*) suspension mixed with distilled water.

To evaluate the performance of mPCR, we tested 181 serotype known isolates (n=45, serotype 1; n=79, serotype 2; n=49, serotype 11; n=8, other serotypes). Compared to the agglutination typing method, the coincidence rates of mPCR for serotypes 1, 2 and 11 were 93.33% (42/45), 97.47% (77/79) and 100% (49/49), respectively. An excellent agreement was found between the mPCR and the agglutination method, with kappa index 0.96±0.03 at the 95% confidence level (*p*-value < 2.2e-16).

### Identification and agglutination characterization of *R. anatipestifer* CH-2Δ *wzy*

The *wzy* of *R. anatipestifer* CH-2 was knocked out by allelic exchange, and the mutant CH-2 *wzy* was identified by PCR(Figure 1). CH-2Δ *wzy* amplified the 16S rRNA fragment, Spec^R^ cassette fragment, and LSR fragment, but did not amplify the *wzy* fragment. All amplicons were confirmed by Sanger sequencing. After continuous culture for 30 generations, the genetic stability of the CH-2Δ *wzy* mutant was confirmed by the same PCR test. A standard antisera slide agglutination test showed that CH-2Δ*wzy* could not agglutinate with the antisera of serotype 2 (Figure 13).

## Discussion

In the present study, we used Pan-GWAS and identified the genetic loci significantly associated with *R. anatipestifer* serotype 1, serotype 2, serotype 10 and serotype 11. Further functional analysis of the loci suggested that these genes are responsible for the synthesis of O-antigen-related polysaccharides. This is consistent with previous studies showing that each serotype of gram-negative bacterial species corresponds to a specific cluster of O-antigen synthesis genes(Aydanian et al., 2011; Kenyon et al., 2017; Liu et al., 2014; Liu et al., 2008; Seif et al., 2019; Wang et al., 2017b). Wang et al. mutated the *AS87_04050* gene (strain Yb2, serotype 2) located in the abovementioned O-antigen gene cluster. The results showed that compared with the wild-type strain, the mutant LPS was defective and lost it’s agglutination ability to serotype 2-positive antisera(Wang et al., 2014). Similarly, Zou et al. mutated the *M949_1603* gene in the above gene cluster in *R. anatipestifer* CH-3 and found that the LPS of the mutant strain lacks the O-antigen chain(Zou et al., 2015). Two other studies reported similar results, that is, the expression of *M949_RS07580* was significantly downregulated in the CH-3 mutant strain, which lacks O-antigen repeat units(Dou et al., 2017; Dou et al., 2018). *M949_RS07580* is also located in the O-antigen gene cluster.

Based on the results of the correlation study between serotype and genome, we predicted and analysed the O-antigen gene cluster of *R. anatipestifer*. The existence of *wzx* and *wzy* means that the O-antigen polysaccharide units are assembled via the Wzx/Wzy-dependent pathway. Functional annotation predicted that the *G148_RS04365* gene of *R. anatipestifer* CH-2 was *wzy* (coding O-antigen polymerase, Wzy; Table 3). *G148_RS04365* protein contains contain 10 transmembrane regions, which is the basic characteristic of Wzy. Furthermore, the *wzy* gene (G148_RS04365) was deleted from *R. anatipestifer* CH-2, and the mutant CH-2Δ agglutinate with antisera of serotype 2, which indicates that the mutant strain antigen was defective. The same phenomenon occurred in Yb2 when the *AS87_04050* gene (predicted nucleoside-diphosphate-sugar epimerase) was knocked out(Wang et al., 2014).The above results also indicate that the O-antigen gene cluster our proposed is closely related to the serotype of *R. anatipestifer*.

In this study, we performed a conservative analysis of the O-AGCs of *R. anatipestifer* in *Flavobacterium* species. It is noteworthy that a similar genetic locus is harboured in some species of *Chryseobacterium* and *Elizabethkingia* (Figure 9).

However, there have been no reports about the O-antigen gene cluster of *Chryseobacterium* and *Elizabethkingia*. Despite this limitation, we found several genes related to O-antigen synthesis in these regions, such as *wbpA*, *wbpD*, *wbpE*, *lptB* and the ABC transporter ATP-binding protein gene(Luo et al., 2017; Shoji et al., 2014). Therefore, for some species of *Chryseobacterium* and *Elizabethkingia*, the abovementioned genomic region may also be the locus of the O-antigen gene cluster.

The comparison of the O-AGCs of a total of 11 serotypes indicates that the O-AGC of *R. anatipestifer* is conserved at both ends and variable in the middle region (Figure 8). Similar phenomena also appeared in *Plesiomonas shigelloides*(Xi et al., 2019), *Escherichia albertii*(Wang et al., 2017b), and *Yersinia pseudotuberculosis*(Kenyon et al., 2017). Variations in the O-AGCs often mean differences in the O-antigen oligosaccharide unit and affect the serological phenotype. Our analysis of the phylogenetic relationship of the O-AGCs from *R. anatipestifer*’s 11 serotypes (38 strains) reveals correspondence between the O-AGCs and their serotypes. In the present study, mPCR based on specific sequence for each serotype to detect three main serotypes was developed. To evaluate the performance of the mPCR method, we compared it with standard slide agglutination serotyping, and the results show that agreement between our method and conventional agglutination methods was very high (kappa = 0.96±0.03), which indicates that the mPCR method could be an alternative to the traditional method. Since first using slide agglutination test to determine serotypes of *R. anatipestifer*(Bisgaard, 1982), no new molecular serotyping scheme has been proposed to conveniently detect the serotype. Therefore, our findings and proposed method were significant for the establishment of system serotyping schemes for *R. anatipestifer*.

## Conclusion

In this work, we revealed that the serotype of *R. anatipestifer* is related to the putative O-antigen gene cluster through a genome-wide association studies and construction of a gene knockout strain. We characterized the O-antigen gene clusters of 11 of *R. anatipestifer* serotypes and demonstrated their genetic diversity. The serotyping mPCR approach defined here will facilitate work on the epidemiological surveillance of *R. anatipestifer*.

