## Supplementary Figure 1 - 4 for "Genetic loci of the *R. anatipestifer* serotype discovered by Pan-GWAS and its application for the development of a multiplex PCR serotyping method"

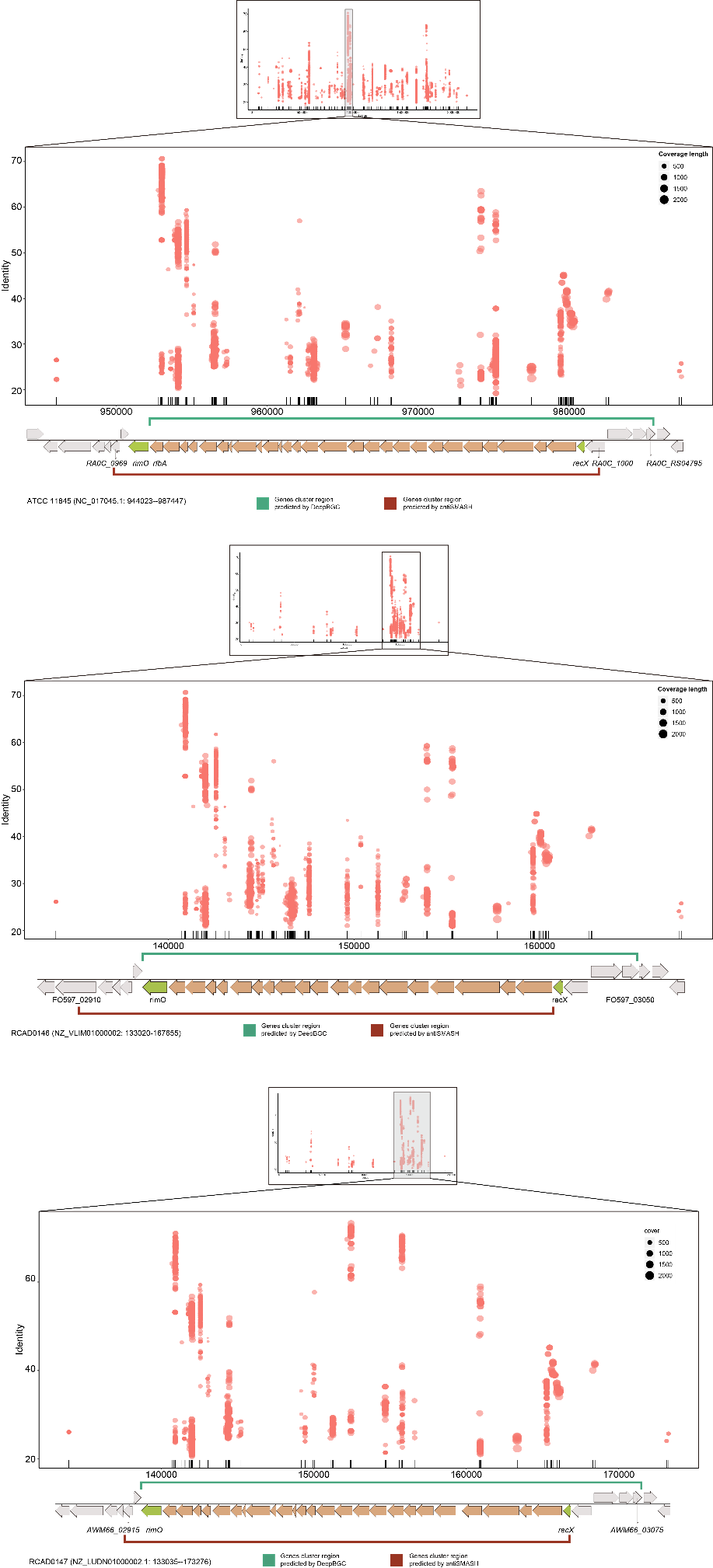

**Supplementary Figure 1.** *R. anatipestifer* serotype 1 (ATCC 11845), 10 (RCAD0146) and 11 (RCAD0147) O-antigen gene cluster location and boundary

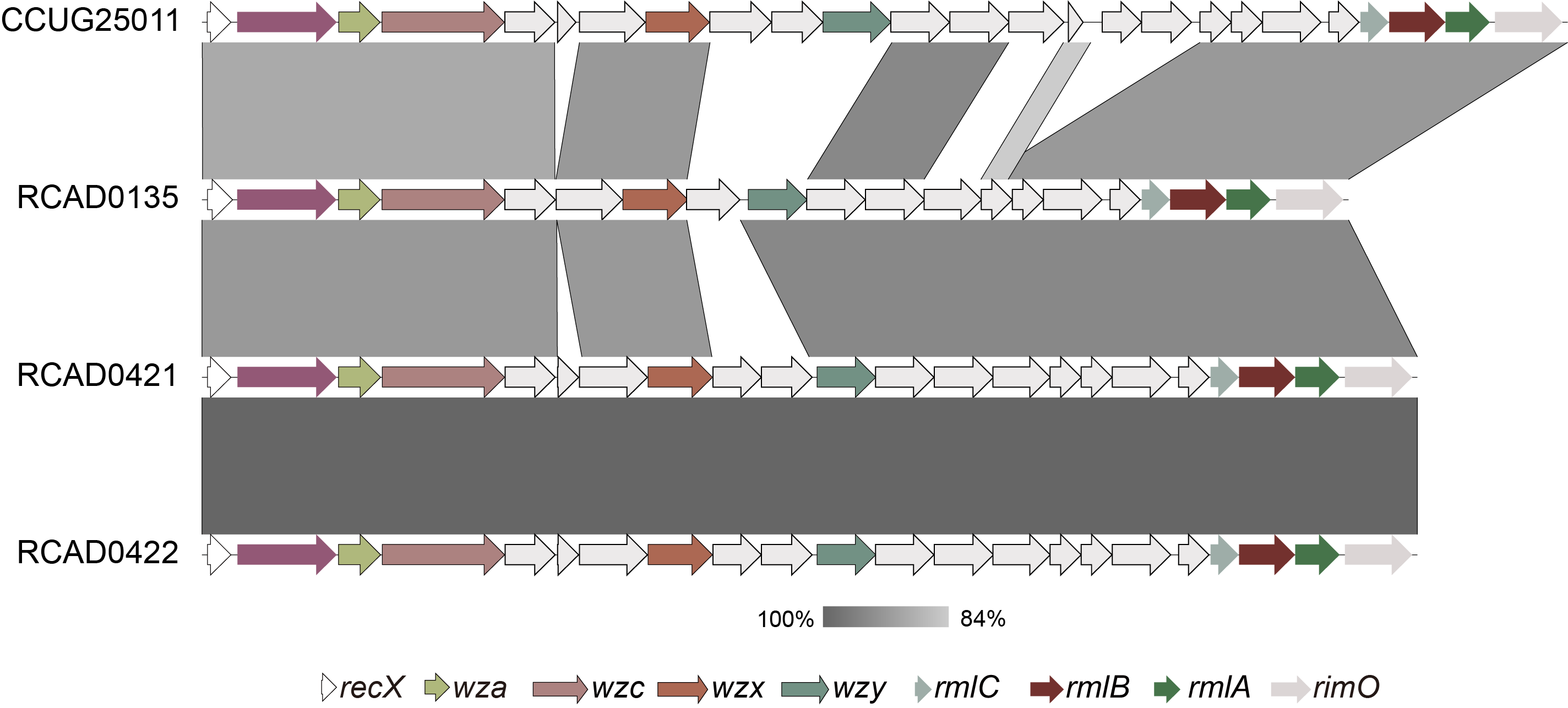

**Supplementary Figure 2.** Comparison of O-antigen gene cluster between serotypes 4 (CCUG 25011)

and serotypes 7 (RCAD0135, RCAD0421 and RCAD0422) of *R. anatipestifer*

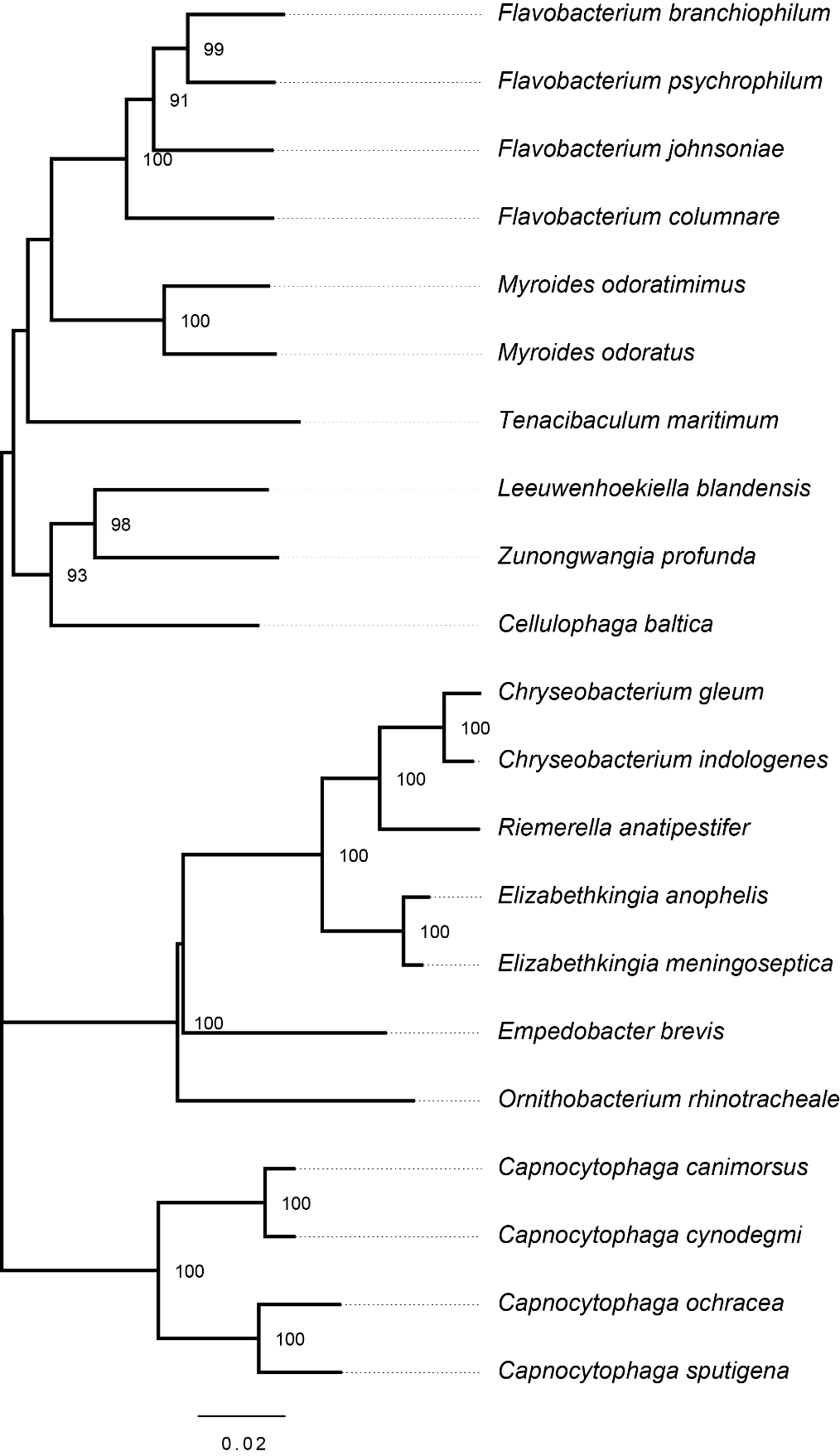

**Supplementary Figure 3.** 16S rDNA NJ (neighbor joining) phylogenetic tree of closely related

species for *R. anatipestifer*. 16S rDNA nucleotide sequence alignment was performed using MAFFT and tree was reconstructed by MEGA 7 with default parameters and 1000 bootstrap replicates.

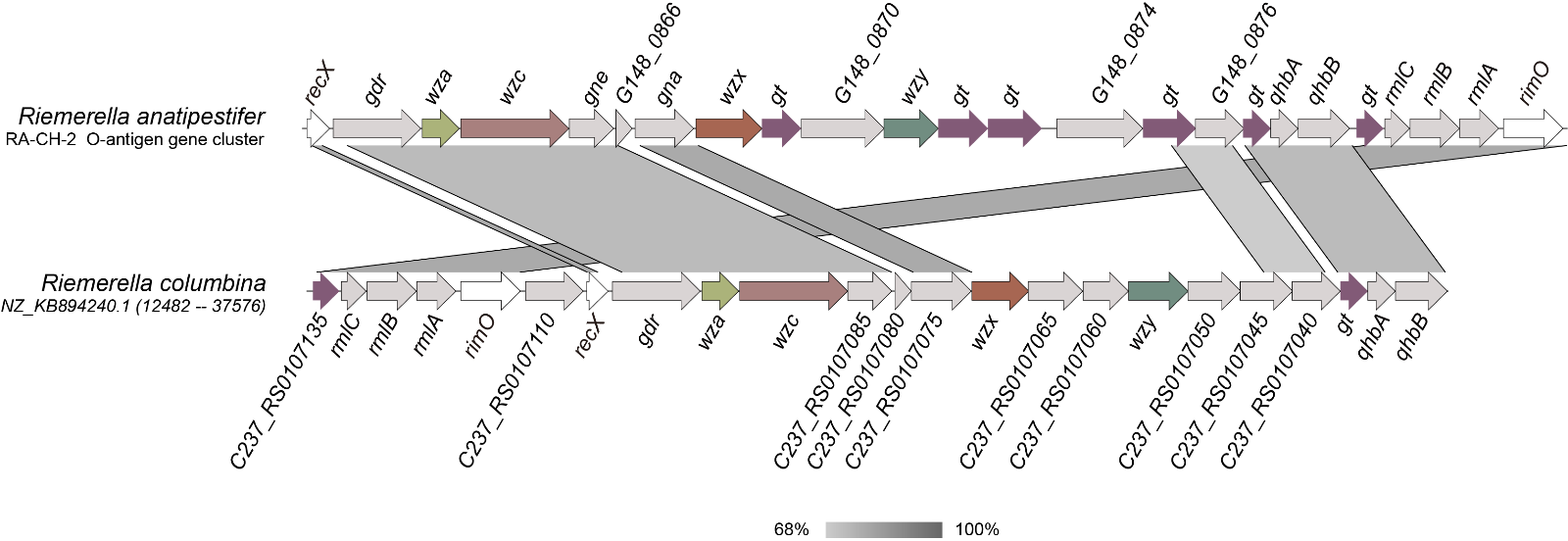

**Supplementary Figure 4.** Comparison between the O-antigen gene cluster of *R. anatipestifer* (CH-2) and the homologous regions of *R. columbina* (DSM 16469)
